# Loss of riboflavin biosynthesis leads to accumulation of select aromatic amino acids and loss of infectivity in *Mycobacterium tuberculosis*

**DOI:** 10.64898/2026.08.04.742562

**Authors:** Neetika Jaisinghani, Arshia Arasappan, Atul Pradhan, Rimanpreet Kaur, Kelin Li, Jeffrey Aubé, Justin R. Cross, Ruben J. Jesus Faustino Ramos, Travis Hartman, Mary L. Previti, Charles K. Vorkas, Jessica C. Seeliger

## Abstract

Riboflavin biosynthesis is required for *in vitro* survival of the human pathogen *Mycobacterium tuberculosis* (*Mtb*). However, despite the lack of a known transporter, growth can be rescued by exogenous riboflavin. The riboflavin biosynthesis pathway is also predicted essential *in vivo*, but whether riboflavin levels available in the host can support survival has not been directly tested. Here we constructed a set of inducible CRISPR interference (CRISPRi) knockdown and targeted gene deletion strains for known riboflavin biosynthesis genes (*ribA2, ribG, ribH, ribC*) as tools to characterize riboflavin requirements, uptake, and metabolite changes and to assess *in vivo* essentiality. We found that riboflavin, but not flavin adenine dinucleotide or flavin mononucleotide, rescued auxotrophy for all strains tested. Further, riboflavin uptake was not affected by disruption of proton motive force or ATP synthesis, suggesting that uptake is not via an active mechanism. Targeted metabolite profiling after removal of riboflavin from growth medium confirmed reduced riboflavin levels. While other riboflavin intermediates were not detected, significant accumulation of aromatic amino acids (Phe, Tyr) was observed across all assayed strains, as well as alteration in a vitamin B9 metabolite. Selecting the *ribC* knockout as a representative strain, we found that riboflavin depletion had a bacteriostatic effect as late as 3 weeks after removal. Unexpectedly, Δ*ribC* lacked infectivity in an aerosol mouse infection, suggesting that the potential to scavenge riboflavin from the host is not sufficient to survive *in vivo*. Overall, our results show that altered metabolism upon loss of riboflavin biosynthesis leads to compromised *Mtb* infectivity.

**IMPORTANCE:** Tuberculosis remains one of the world’s most deadly infectious diseases, underscoring the need to explore new drug targets. Riboflavin (vitamin B2) biosynthesis has emerged as a promising target because Mycobacterium tuberculosis (*Mtb*) depends on this pathway for survival. The riboflavin pathway also produces metabolites that modulate host mucosal-associated invariant T (MAIT) cell activity, towards understanding potential strategies for host-directed therapies. Here we found that disrupting riboflavin biosynthesis led to not only compromised survival, but also widespread changes to metabolism and loss of the ability to establish infection in an animal model. These findings improve our understanding of how *Mtb* adapts to metabolic stress, with implications for developing drugs that target riboflavin biosynthesis and for alterations in host immunity to be explored in future studies.

## INTRODUCTION

Riboflavin (vitamin B2) is synthesized *de novo* by all plants and most bacteria and fungi, but not humans, who must acquire it from their diet. Riboflavin is a precursor to essential coenzymes, flavin adenine dinucleotide (FAD) and flavin mononucleotide (FMN), that are required for most biochemical reactions involved in essential carbohydrate, lipid and protein metabolism. Riboflavin biosynthesis, derived from the purine and pentose phosphate pathways (**Figure 1**), is predicted to be essential for *in vitro* growth of the human pathogen, *Mycobacterium tuberculosis* (*Mtb*), based on genome-wide null mutant screens (1,2). These observations have underscored this pathway as an attractive target for antimycobacterial therapy and motivated the pursuit of inhibitors (3,4).

**Figure 1.**
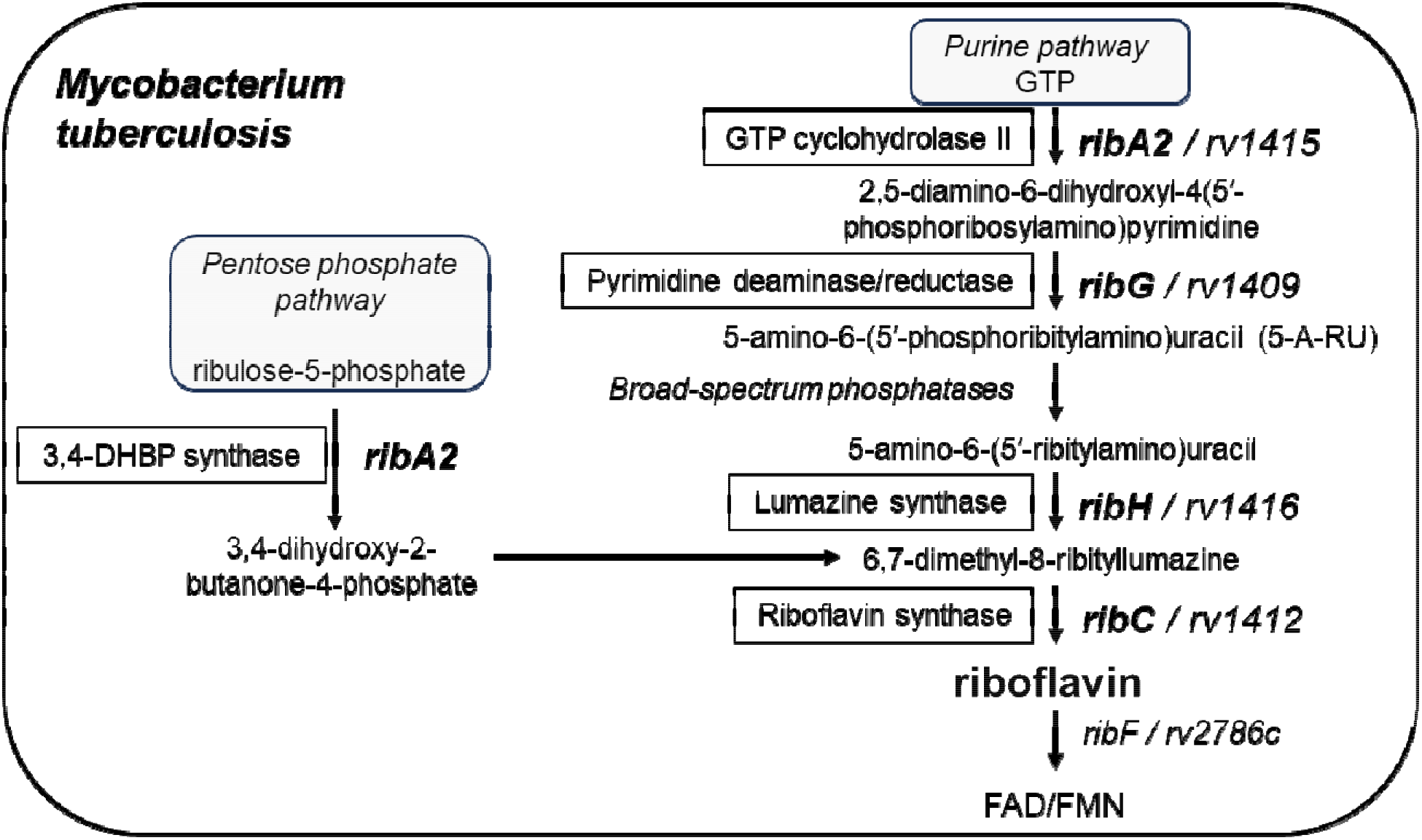
Riboflavin metabolism in *Mtb*. The riboflavin metabolism pathway in *Mtb* uses products from two distinct pathways: purine synthesis and the pentose phosphate pathway (shaded boxes). Gene names are highlighted in bold italics and the class of the encoded enzymes are boxed.

Beyond its metabolic role, the riboflavin biosynthesis pathway is also a source of activating metabolite ligands for mucosal-associated invariant T cells (MAIT), an innate-like T cell subset restricted by the highly conserved mammalian major histocompatibility complex class I-related molecule, MR1(5). MR1 presents vitamin B metabolites, of which one of the most potent, 5-(2-oxopropylideneamino)-6-d-ribitylaminouracil (5-OP-RU), is synthesized from the riboflavin intermediate 5-amino-6-(5′-ribitylamino) uracil (5-A-RU) through a condensation reaction with the glycolytic intermediate methylglyoxal (6,7). In addition to riboflavin biosynthesis pathway intermediates, additional MR1 ligands include those derived from vitamins B9 (folic acid) and B6 (7–9).

The dual significance of riboflavin biosynthesis in mycobacterial survival and host immune recognition provided the rationale for the present study to dissect this pathway through genetic manipulation of virulent *Mtb* and thereby define its role in bacterial physiology and as a potential therapeutic target. Importantly, targeting riboflavin biosynthesis also hinges on the pathogen’s inability to salvage riboflavin from the host and thus highlights the importance of potential mechanisms of riboflavin uptake and directly testing *in vivo* essentiality. *Mtb* is assumed to lack a riboflavin transporter based on the absence of homologues to known bacterial transporters, although this does not rule out the possibility of a novel transporter (10). Indeed, two transposon sequencing (Tnseq) analyses showed that mutants with insertions in riboflavin biosynthesis genes survived on medium supplemented with riboflavin, supporting uptake by *Mtb* (11,12).

Consistent with these findings, we found in the present study that *Mtb* strains lacking the riboflavin biosynthetic machinery were able to grow and survive *in vitro* when micromolar exogenous riboflavin was supplied, further underscoring riboflavin auxotrophy. Two recent studies independently reported similar observations (13,14). Using targeted gene deletion strains, we further characterized the minimum riboflavin concentration required for survival in culture and showed that riboflavin uptake in mycobacteria is not strongly dependent on active transport driven by ATP or proton motive force. We confirmed depletion of intracellular riboflavin across *Mtb* riboflavin auxotroph strains and observed accumulation of aromatic acids, suggesting a significantly altered metabolism. Using Δ*ribC* as a representative auxotroph strain, we demonstrate *Mtb* riboflavin import is insufficient to establish murine aerosol infection. These findings collectively provide indirect but compelling evidence for the absence of a dedicated riboflavin transporter in *Mtb* and underscore the value of riboflavin biosynthesis as a therapeutic target.

## RESULTS

### Inducible CRISRi knockdown of riboflavin biosynthesis genes leads to a growth defect that is rescued by exogenous riboflavin

As endogenous riboflavin synthesis is predicted to be essential for mycobacterial survival *in vivo* (13,14), we initially selected inducible CRISPRi (15) to generate anhydrotetracycline (ATc)-inducible knockdown of the riboflavin biosynthesis genes *ribA2* (*rv1415*), *ribG* (*rv1409*), *ribH* (*rv1416*), and *ribC* (*rv1412*) in *Mtb* H37Rv (**Figure 1, 2A, Supplemental Table S1**). CRISPRi knockdown of *ribA2*, *ribH*, or *ribC* was validated by qRT-PCR, which confirmed significant decreases in target mRNA relative to non-targeting controls (**Figure 2B-D**). Knockdown of the gene *ribG* using a range of protospacer adjacent motifs (PAMs) (**Supplemental Table S2**) was not achieved with this method: None of the constructed strains showed an ATc-dependent decrease in *ribG* transcription (**Supplemental Figure S1**), in contrast to a recent study (13). Thus, further characterization proceeded with only the *ribA2*, *ribH*, and *ribC* CRISPRi strains. We also measured expression of all *rib* and other predicted co-cistronic genes for each strain to assess potential polar effects. In particular, *ribA2, ribH*, and *rv1417* (which does not have a known role in riboflavin biosynthesis) form a predicted operon. Indeed, knockdown of *ribA2* or *ribH* significantly downregulated transcription of downstream co-cistronic genes, whereas *ribC* knockdown did not affect other genes in the pathway (**Figure 2C-D**). These results confirm successful knockdown of target genes, but also indicate that targeted knockdown of *ribA2* or *ribH* is not achieved with CRISPRi, complicating the interpretation of phenotypes and their specific attribution to *ribA2* or *ribH* function.

**Figure 2.**
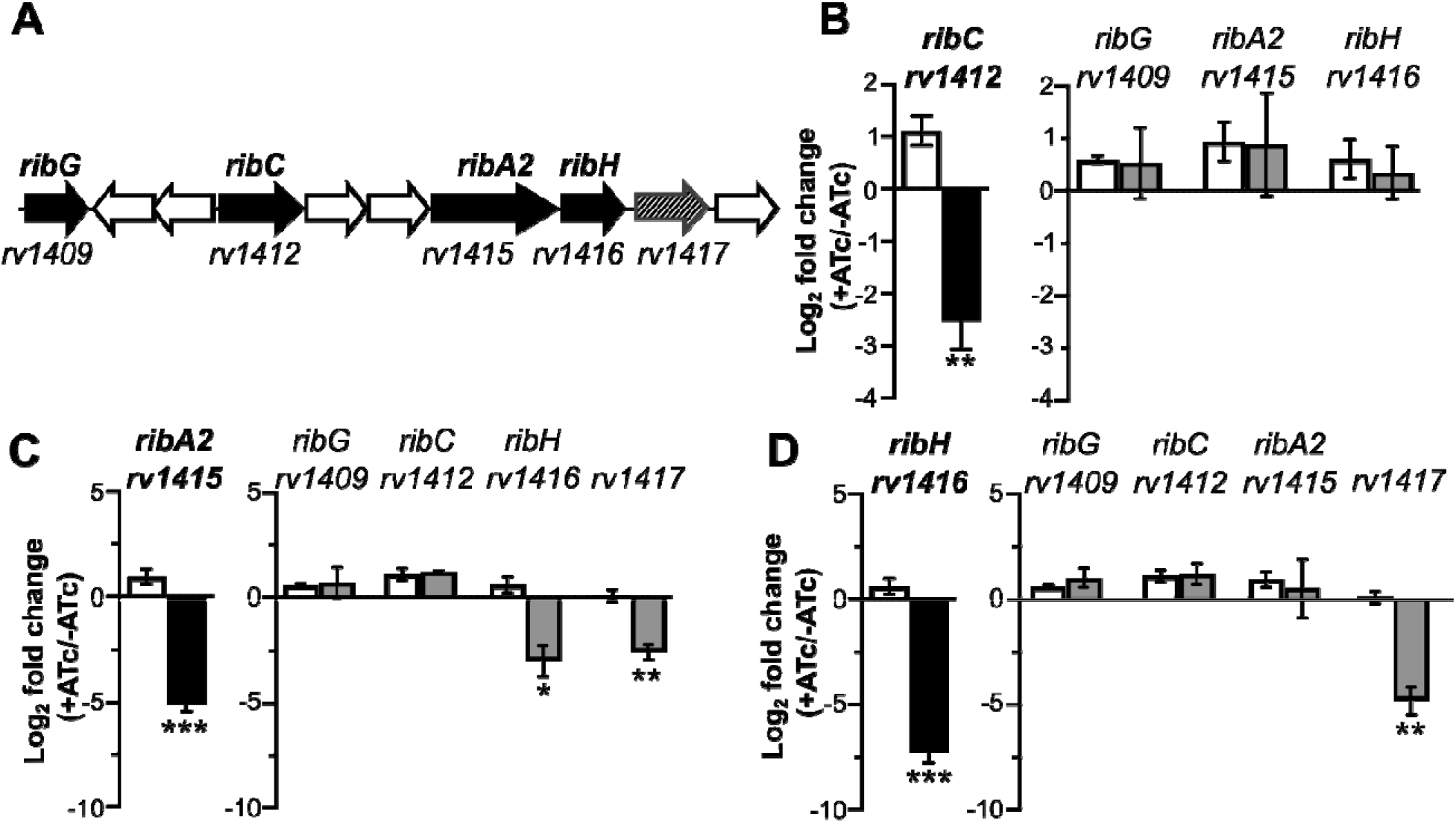
CRISPRi knockdowns strains exhibit reduced expression of respective riboflavin metabolism genes upon ATc induction. **(A)** Locus for genes encoding riboflavin biosynthesis enzymes. The three genes *ribA2, ribH,* and *rv1417* form a predicted operon. Erdman *Mtb* strains encoding sgRNA targeting a non-targeting (NT) control (white bar) or **(B)** *ribC*, **(C)** *ribA2*, or **(D)** *ribH* were cultured with 0 or 100 ng/mL ATc for 3 days. Total RNA extracted from each condition was used to generate cDNA for qRT-PCR. Raw Ct values obtained for respective genes in each condition were normalized to the housekeeping gene *16SrRNA* and the ratio +ATc/-ATc calculated to determine fold change. Data shown are mean ± SEM from 3 independent experiments. Statistical significance was calculated by 2-tailed t-test (* *p* <0.05, ** *p* <0.01, *** *p* <0.001).

To test the effect of *rib* knockdown on mycobacterial survival *in vitro*, we then measured bacterial growth in liquid culture in the presence and absence of ATc. The growth of all knockdown strains was attenuated in the presence of ATc as compared to untreated or non-targeting controls, with *ribA2* knockdown generating the largest growth defect (**Figure 3A**).

**Figure 3.**
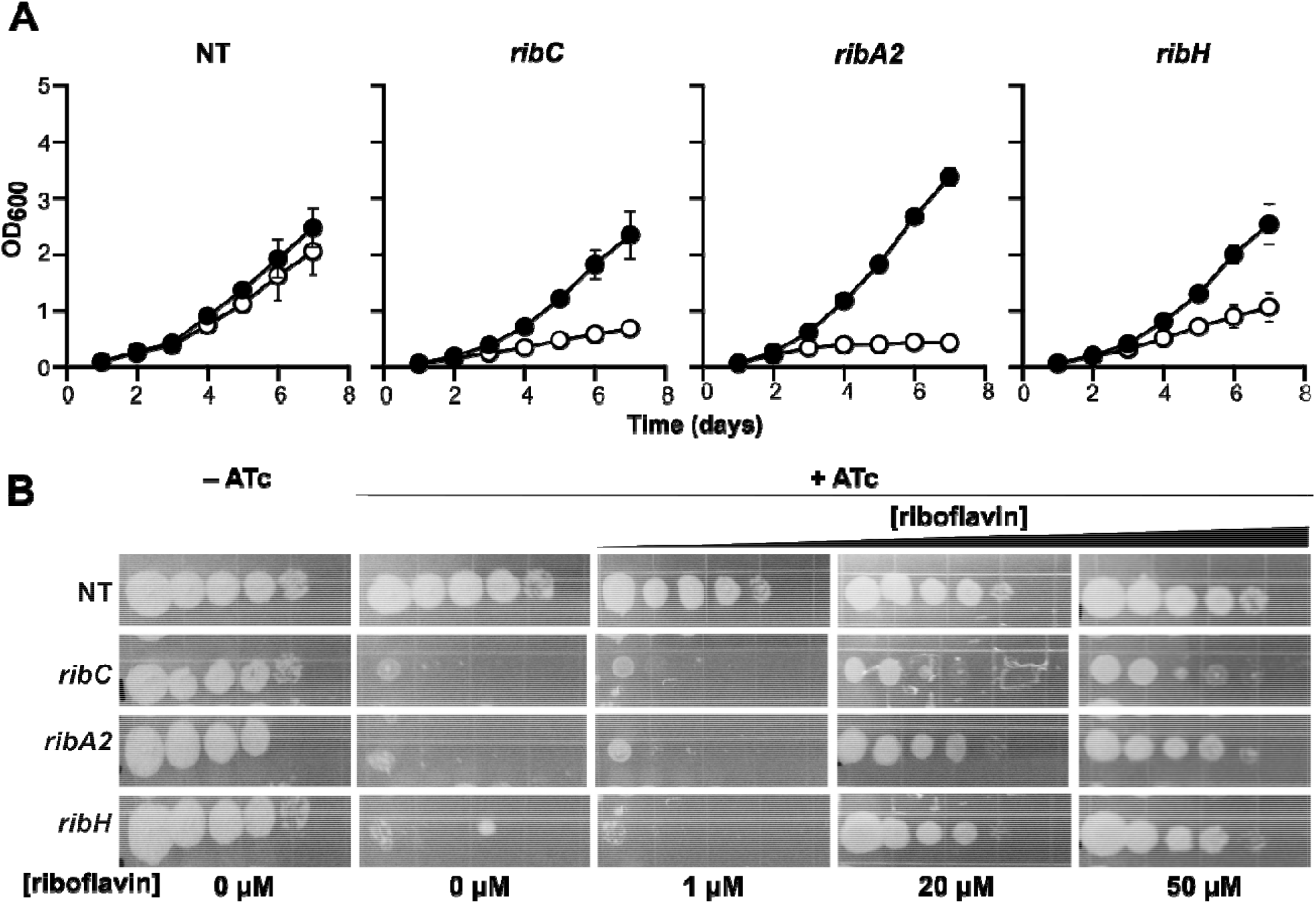
Growth attenuation upon CRISPRi knockdown of riboflavin biosynthesis genes is rescued by exogenous riboflavin. Erdman *Mtb* strains encoding sgRNA targeting a non-targeting (NT) control, *ribC, ribA2*, or *ribH* were grown to log phase and then **(A)** sub-cultured to OD_600_ 0.015 in growth medium without ATc (black) or with 100 ng/mL ATc (white) or **(B)** diluted to 10^5^ cells/ml and spotted as 10-fold serial dilutions on agar without ATc or with 100 ng/mL ATc and riboflavin at 0, 1, 20 or 50 µM. Plates were imaged after incubating for 12-15 d. Data shown are (A) the mean of 3 independent experiments ± SEM or (B) from a single experiment.

To confirm growth rescue with exogenous riboflavin as previously observed by Chengalroyen *et al*. (14), log-phase liquid cultures of each CRISPRi strain were spotted on agar with or without ATc along with various concentrations of riboflavin. All tested *rib* knockdown strains failed to grow appreciably in the presence of ATc without riboflavin supplementation compared to uninduced and non-targeting controls (**Figure 3B**). However, supplementation with 20 µM or 50 µM, but not 1 µM, riboflavin rescued growth. These results further confirm that *Mtb* has the capacity to uptake and assimilate exogenous riboflavin to survive *in vitro*.

### Genetic deletion of essential rib genes using exogenous riboflavin supplementation

Inducible CRISPRi technology, while convenient and useful for investigation of this essential pathway, ultimately posed various challenges. First, we were unable to achieve successful *ribG* knockdown despite testing six different PAM oligonucleotides. Second, CRISPRi knockdown had epistatic effects that suppressed expression of downstream co-cistronic genes (**Figure 2**), complicating interpretation and attribution of phenotypes to specific gene functions.

Third, we observed that strains cultured without kanamycin regained the ability to grow without riboflavin, likely due to loss of the plasmid encoding the suppressing PAM oligonucleotide, when under the selective pressure of downregulating an essential gene (**Supplemental Figure S2**). Thus, selection with kanamycin is absolutely necessary to maintain CRISPRi knockdown, but its toxicity to eukaryotic cells and animals during extended use limits the utility of the current CRISPRi system in infection models. Finally, while significant suppression of expression and growth was observed across strains upon CRISPRi knockdown, extraction and targeted metabolomic analysis by LC-MS/MS indicated that only *ribA2* knockdown significantly depleted riboflavin after 3 d culturing with ATc induction (**Supplemental Figure S3**). This suggests that riboflavin utilization by *Mtb* does not efficiently deplete stores over the tested period of depletion and under conditions of *rib* knockdown used for metabolite extraction.

Nevertheless, the CRISPRi knockdown strains enabled validation on solid growth medium that exogenous riboflavin rescues auxotrophy, which led us to generate *rib* null strains as an accessible alternative. Using recombineering followed by recovery on riboflavin-containing medium, we were unable in multiple attempts to recover legitimate recombinants for any *rib* target in the *Mtb* Erdman strain. However, we successfully obtained Δ*ribC* and Δ*ribG* strains in H37Rv, which we proceeded to use as the parent strain for all targeted deletions. Since we had reason to believe from CRISPRi knockdown that *ribA2*, *ribH* and *rv1417* are co-transcribed, we chose to address potential epistatic effects by first generating the operonic deletion Δ*ribA2-ribH-rv1417* and then complementing with the appropriate genes driven by a native promoter (1000 nt upstream of *ribA2*). For simplicity, we herein refer to these strains as Δ*ribA2* (Δ*ribA2-ribH-rv1417:: ribH-rv1417*) and Δ*ribH* (Δ*ribA2-ribH-rv1417:: ribA2-rv1417*) with a single complement strain for both (Δ*ribA2-ribH-rv1417:: ribA2-ribH-rv1417*) (see **Supplemental Table S1**). All deletions were confirmed by sequencing the targeted locus and measuring expression of *rib* genes and *rv1417* by qRT-PCR to confirm loss/retention of expression, as relevant (**Supplemental Figure S4**). Based on RNA expression, the partial complementation strategy was successful for Δ*ribH* and Δ*rv1417* and the operon complement restored expression of all three genes. In contrast, Δ*ribA2* displayed reduced transcription of *ribH* and *rv1417* similar to that observed for the *ribA2* CRISPRi knockdown (**Figure 2C, Supplemental Figure S4**). This suggests that there are regulatory elements within the coding sequence of *ribA2* that affect *ribH* and *rv1417* transcription. Aside from Δ*ribA2*, deletion with partial complementation was more specific to the targeted gene than CRISPRi-mediated knockdown, so we pursued further characterization with this set of strains.

Consistent with the CRISPRi knockdown strains, *Mtb rib* deletion strains failed to grow in the absence of riboflavin both in liquid culture and on solid medium (**Figure 4A-F**). This included loss of function in *ribG,* which we had been unable to test by CRISPRi knockdown. The growth of these strains in the presence of riboflavin was comparable to that of the wild type and respective complements. Exogenous riboflavin supplementation itself had no effect on the growth of wild type and complemented strains (**Supplemental Figure S5**). Since riboflavin is the sole precursor for FAD and FMN and these metabolites are coenzymes for many essential cellular processes, we also tested FAD and FMN supplementation, but supplementation up to 20 µM did not rescue growth in *Mtb rib* mutants (**Supplemental Figure S6**), consistent with lower concentrations tested in a previous report (13).

**Figure 4.**
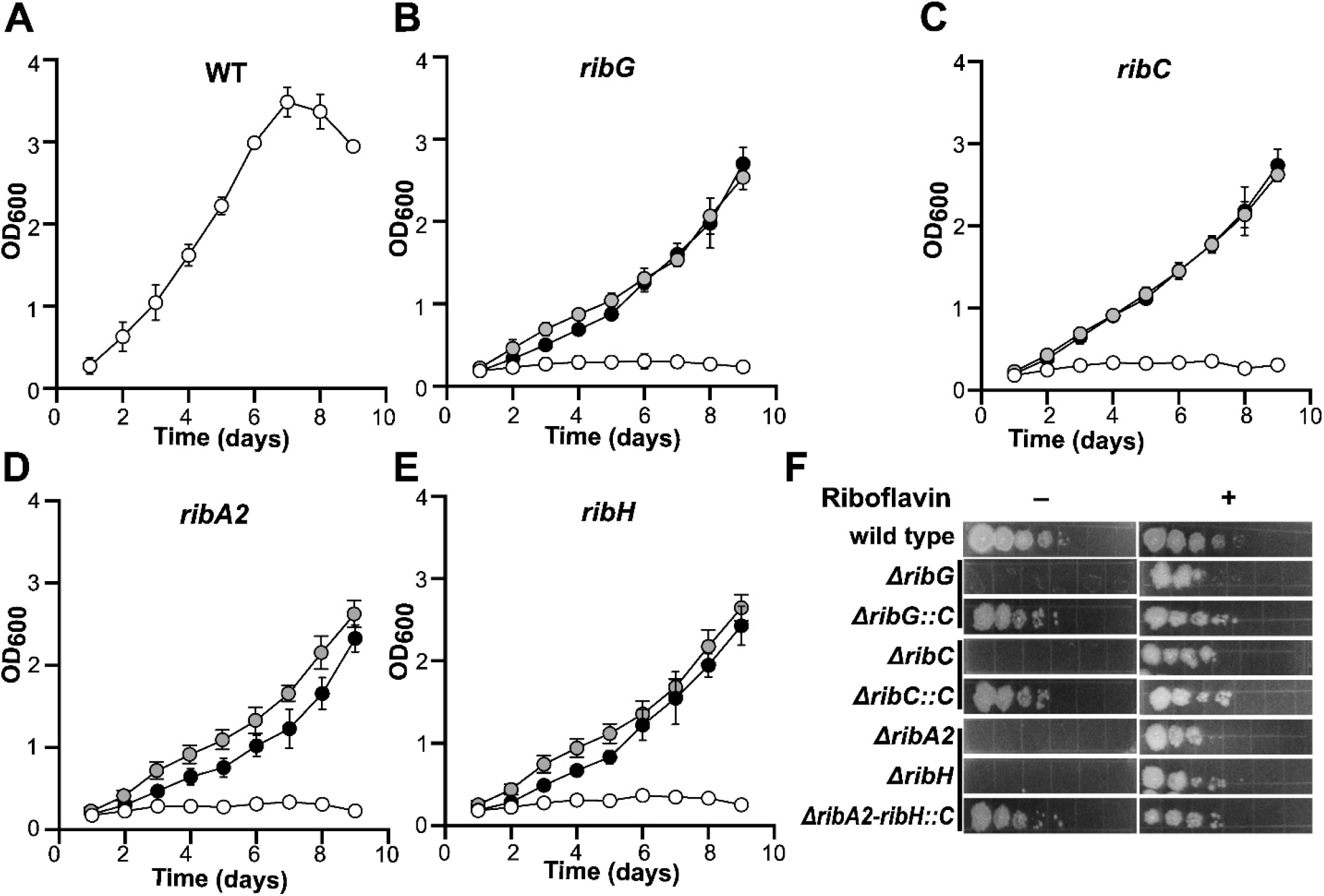
Targeted *rib* gene knockout confirms essentiality of riboflavin auxotrophy and rescue by exogenous riboflavin. Growth of *Mtb* H37Rv **(A)** wild type, **(B)** Δ*ribG,* (C) Δ*ribC*, **(D)** Δ*ribA2*, and **(E)** Δ*ribH* as measured by OD_600_ daily for 9 d without riboflavin (white circles) or with 20 µM riboflavin (black circles). Data for corresponding complement strains are also shown (grey circles). The complement strain in (E) is Δ*ribA2-ribH-rv1417::ribA2-ribH-rv1417* and thus is also the appropriate complement for Δ*ribA2* in (D) (see full strain genotypes in **Supplemental Table S1**). Data shown are mean ± SEM from 3 independent experiments. **(F)** Strains grown as in (A)-(E) were diluted to 10^5^ cells/ml and spotted in serial 10-fold dilution on agar with and without 20 µM riboflavin. Data shown are representative of 2 independent experiments.

### Riboflavin uptake in mycobacteria is unaffected by disruption of the proton gradient or ATP synthesis

While a riboflavin importer has been described in Corynebacteria (16), Mycobacteria do not encode a homologue of any known riboflavin transporter, leading to the inference that *Mtb* cannot take up riboflavin. However, since both we and others demonstrated riboflavin auxotrophy (13,14), we next investigated possible mechanisms of riboflavin uptake in *Mtb*. We confirmed that wild type *Mtb* is able to take up riboflavin, as determined from scintillation counts associated with total lysates, with maximum observed levels achieved at 60 min post incubation with 2 μM ^3^H riboflavin (**Figure 5A**).

**Figure 5.**
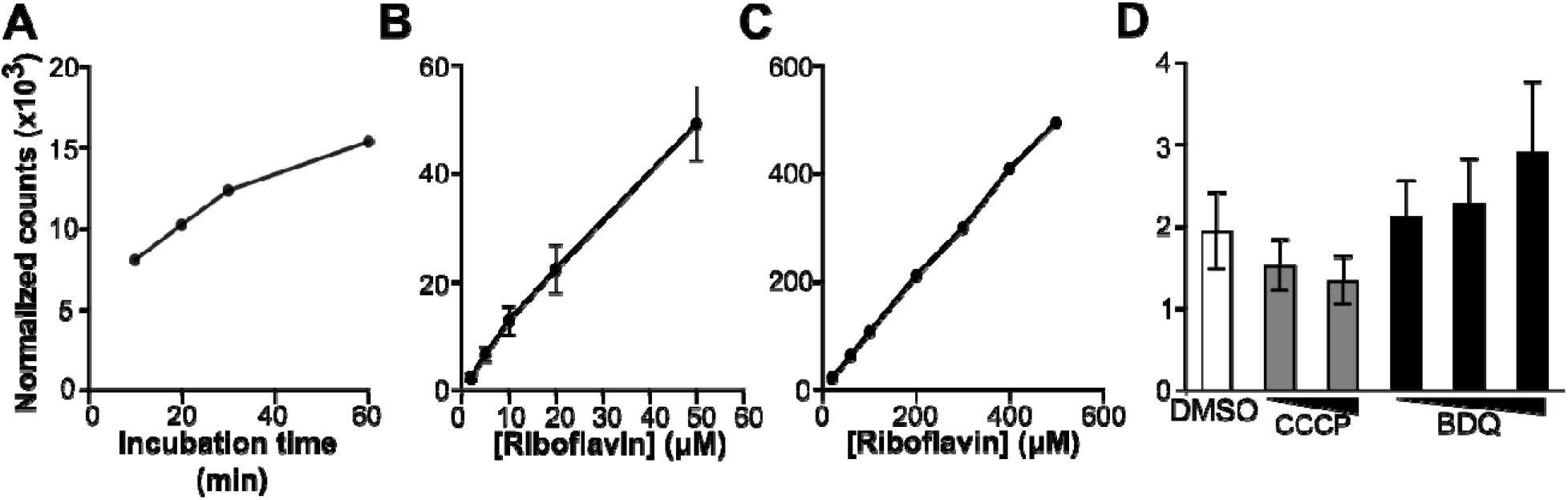
Riboflavin uptake is directly proportional to riboflavin concentration and largely unaffected by disruption of the proton gradient or ATP synthesis. *Mtb* H37Rv was incubated **(A)** with 2 µM ^3^H riboflavin for the indicated times prior to harvesting, lysis, and scintillation counting, as a measure of intracellular ^3^H riboflavin; **(B)** with exogenous riboflavin at the indicated total concentrations, with ^3^H riboflavin included at 2 µM. Data are mean ± SEM from three independent experiments. **(C)** Same as in (B), except that ^3^H riboflavin was included at 20 µM. **(D)** *Mtb* H37Rv was treated for 30 min with DMSO or CCCP (at 20 µM or 200 µM) or bedaquiline (BDQ, at 3.6 µM, 36 µM or 360 µM) followed by incubation with 2 µM ^3^H riboflavin for 1 h. Data shown are mean ± SEM from 3 independent experiments.

We then assessed the dependence of uptake on riboflavin concentration across a range that spans concentrations sufficient to support *in vitro* growth (2 to 50 µM). Across this range, the normalized ^3^H riboflavin signal in the lysate was consistently directly proportional to riboflavin concentration (**Figure 5B**). We also evaluated concentrations up to 500 mM and obtained similar results (**Figure 5C**). We next examined whether riboflavin uptake was sensitive to perturbation of energy-dependent processes. We measured ^3^H riboflavin uptake in *Mtb* cells after treatment with the proton-pump inhibitor carbonyl cyanide m-chlorphenyl hydrazone (CCCP) or ATP synthase inhibitor bedaquiline (F0/F1 ATPase inhibitor; BDQ). Neither CCCP nor BDQ significantly affected riboflavin uptake (**Figure 5D**). Although CCCP induced a dose-dependent decrease in uptake, this effect may rather be related to effects on cell viability (17). Together, these results support that *Mtb* riboflavin transport does not occur via active transport that relies on ATP or proton motive force.

### Targeted deletion of Mtb rib genes results in riboflavin depletion and accumulation of phenylalanine and tyrosine

To assess how riboflavin biosynthesis disruption perturbs *Mtb* metabolism, we performed targeted LC-MS of cell extracts derived from *Mtb* H37Rv WT and *ribA2*, *ribG*, *ribH*, and *ribC* KO strains. *Mtb* strains were grown initially in aerating culture with riboflavin and then transferred to filters on liquid medium (18) without riboflavin for 72 hours. As expected, riboflavin concentrations were significantly decreased in cell extracts of all *Mtb rib* KO strains relative to WT, supporting that all strains are incapable of de novo riboflavin biosynthesis (**Figure 6A**). No other metabolites from the riboflavin pathway were detected from either wild-type or knockout strain samples, as discussed further below. However, from the targeted panel (see **Supplemental Table S5** for full list), we observed accumulation of the aromatic amino acids, phenylalanine and tyrosine in the *Mtb rib* knockout strains compared to the wild type. Phenylalanine (**Figure 6B**) and tyrosine (**Figure 6C**) were significantly increased in cell extracts. Consistent with this observation, we also detected increased concentration of the phenylalanine metabolite phenylethylamine (**Figure 6D**). On the other hand, tryptophan showed statistically significant, but only 10-20%, increases across strains (**Figure 6E**) and further analysis of tryptophan metabolites did not show statistically significant changes or consistent trends (**Supplemental Table S5**). Taken together, our data demonstrate a consistent metabolic phenotype upon disruption of *Mtb* riboflavin biosynthesis that includes accumulation of two aromatic amino acids (Phe, Tyr) and the related metabolite across *rib* knockout strains.

**Figure 6.**
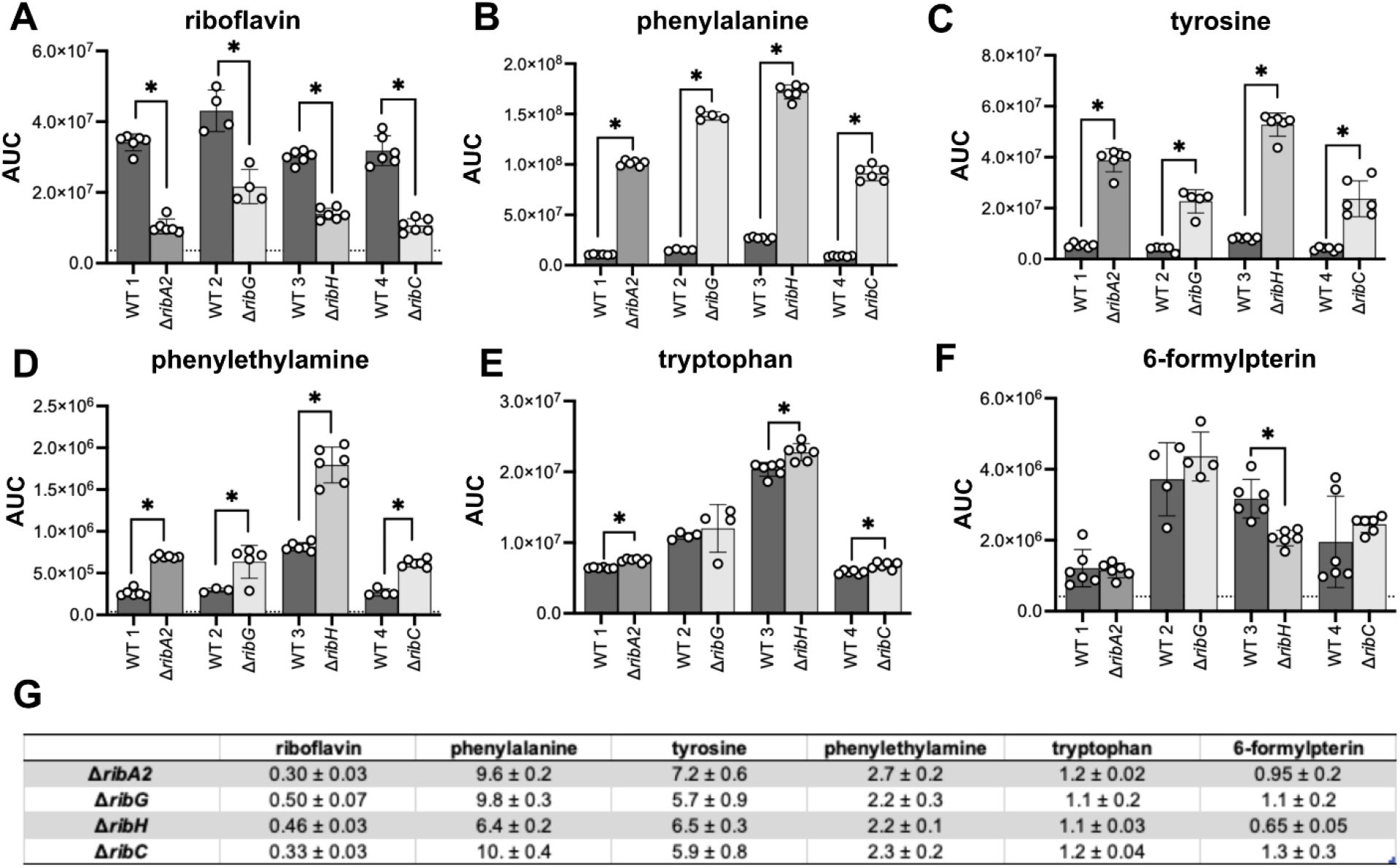
Riboflavin is depleted and phenylalanine and tyrosine accumulate in *rib* gene deletion mutants cultured without riboflavin. (**A)** Riboflavin, **(B)** phenylalanine, **(C)** tyrosine, **(D)** phenylethylamine, **(E)** tryptophan, and **(F)** 6-formylpterin were quantified by LC-MS from extracts of *Mtb* H37Rv *rib* knockout strains (Δ*ribA2*, Δ*ribG*, Δ*ribH*, Δ*ribC*) and the wild type (WT) after culturing for 3 d on medium without riboflavin. Data are the ratio of average AUC ± SEM for n= 4-6 biological samples. Samples for each null strain and a wild-type comparator were prepared together in one independent experiment. The dotted line represents the average value of background signal based on extraction performed without bacteria. Statistical significance was determined using unpaired Student’s t-test (\**p* <0.05). **(G)** Table of fold changes (mutant/wild type) for the metabolites shown in (A-F).

We further examined not only riboflavin intermediates, but also various classes of vitamin B metabolites as other sources of MAIT ligands that were part of the targeted metabolite panel. As noted above, no riboflavin intermediates were detected above background, including the MR1 ligand precursor 5-A-RU and MR1 ligands 5-OP-RU and 6,7-dimethyl-8-ribityllumazine (DMRL), likely due to their instability in solution (14,19). Metabolites from folate (vitamin B9) biosynthesis are inhibitory ligands of MR1 (7). They are also related to aromatic amino acid biosynthesis through shared use of chorismate, which is transformed into prephenate to make Phe/Tyr or into para-aminobenzoic acid (PABA) and coupled to pterin to form folate. While acetyl-6-formylpterin (Ac-6FP) was not detected, another folate breakdown product, 6-formylpterin (6-FP), was moderately decreased in Δ*ribH* but not in other knockout strains (**Figure 6F**). Other MR1 ligands such as those related to other B vitamins (pyridoxal-5’-phosphate, pyridoxamine, nicotinic acid) could not be distinguished by the profiling method used. Nevertheless, our results indicate that disruption of *Mtb* riboflavin biosynthesis impacts *Mtb* metabolism beyond vitamin B2.

### ΔribC survives in vitro following riboflavin depletion, but fails to establish murine aerosol infection

Given the consistent metabolic response to loss of riboflavin biosynthesis across all knockout strains, we chose Δ*ribC* as a representative strain for further characterization. Despite the observed decrease in riboflavin levels by LC-MS, removal of riboflavin had only a moderate effect on Δ*ribC* survival even after 21 days of depletion (**Figure 7A**), indicating that remaining levels of riboflavin were sufficient to support survival *in vitro*.

**Figure 7.**
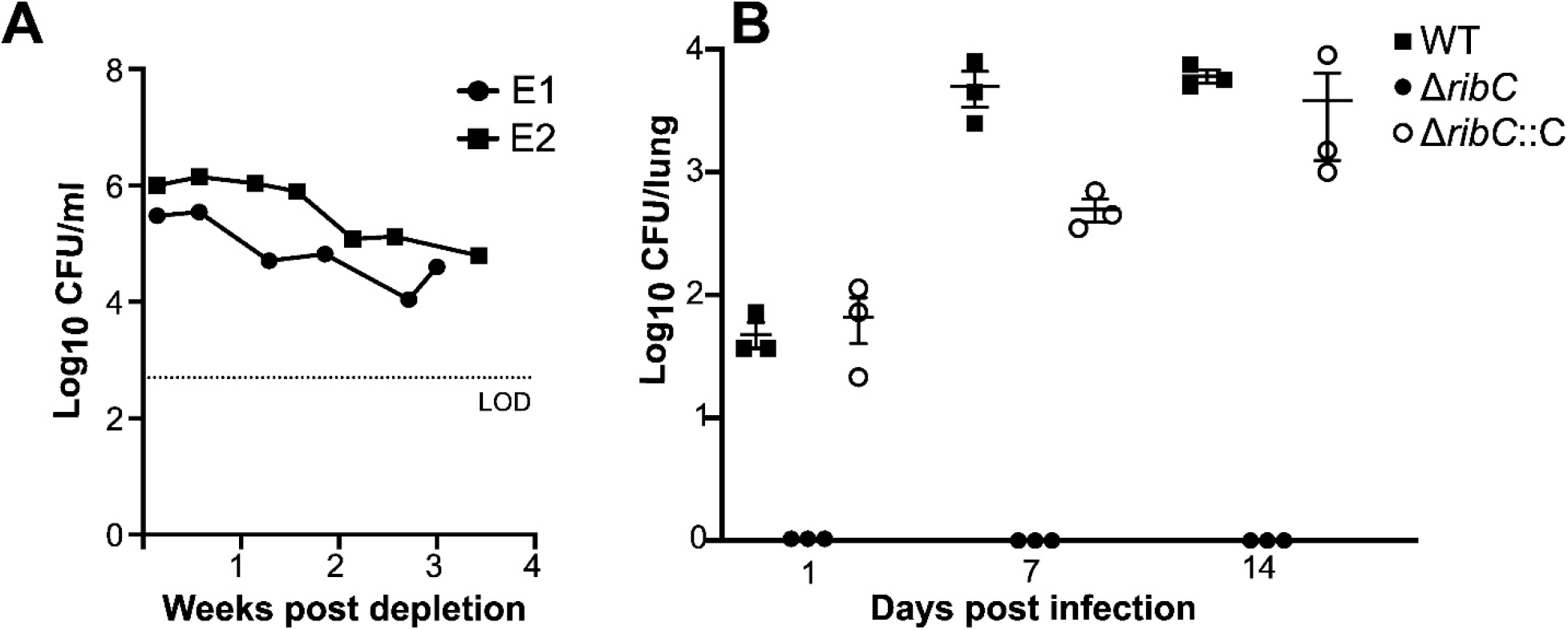
Δ*ribC* survives without exogenous riboflavin *in vitro*, but does not establish infection via the aerosol route in mice. **(A)** *Mtb* H37Rv Δ*ribC* grown to log phase with 20 µM riboflavin was subcultured to OD_600_ 0.01 into medium without riboflavin at day 0. Survival was monitored by CFU enumeration on agar with 20 µM riboflavin. Data from 2 independent experiments (E1, E2) are shown. The dotted line represents limit of detection (LOD). **(B)** C57BL/6 mice were infected with aerosolized Δ*ribC* and CFU/lung were enumerated at the indicated times post infection. Data shown are mean ± SEM CFU/lung (n = 3 per sample group) and are representative of 2 independent experiments.

We next tested if riboflavin auxotrophy could be complemented by the host by measuring Δ*ribC* survival after murine infection. We performed aerosol infection of C57BL/6 mice using a low dose (50-100 CFU/lung) and compared growth to the wild type and complement strains. We found that Δ*ribC* could not be recovered from infected lungs on agar with riboflavin even as early as one day post infection, indicating an infectivity defect that could be complemented (**Figure 7B**).

## DISCUSSION

Previous work has predicted the essentiality of *rib* genes and two studies in defined medium containing riboflavin further indicated that auxotrophy can be rescued exogenously (11,12). Most recently, two additional studies from Chengalroyen et al. used CRISPRi knockdown and targeted gene knockout to confirm this prediction (13,14). Our current report builds on these results, with important additions. First, we did not observe ATc-dependent *ribG* knockdown in any of six *ribG*-targeted CRISPRi strains that we tested (**Supplemental Figure S1**). Chengalroyen *et al.* 2024 reported riboflavin auxotrophy in an analogous CRISPRi *ribG* strain, but the PCR data did not indicate depletion of *ribG* transcripts, similar to our results (see Figure 2 in Chegalroyen et al. 2024). This indicates a contradiction in the data and a possible discordance in the strain reported as a *ribG* knockdown in different assays. We were thus motivated to pursue instead the targeted gene knockout Δ*ribG*, with which we confirmed riboflavin auxotrophy.

Second, we more fully characterized the effect of knocking down or knocking out *ribA2* or *ribH* by checking expression of not only the target, but also the other *rib* loci and genes in the operon (**Figure 2A, Supplemental Figure S4**). The downregulation of downstream genes underscored the importance of accounting for epistatic effects in assigning phenotypes and function to individual *rib* genes, and we took this into account in the design of our knockout strains (**Supplemental Table S1**). Finally, Chegalroyen et al. 2024 tested FAD and FMN complementation at 2 µM, well below the concentration used for riboflavin complementation. We here further confirmed the lack of FAD and FMN complementation up to 20 µM. This result may be expected, as FAD and FMN carry explicit charges and are unlikely to cross the membrane in the absence of a dedicated transporter.

Related to this, we followed up on riboflavin complementation to explore the mechanism of uptake. Riboflavin transporters have been identified in other bacteria; of these, the most closely related are *Streptomyces davawensis* (20) and *Corynebacterium glutamicum* (16), which encode RibM/PnuX. However, *Mtb* and other mycobacteria do not encode homologues of this or any other known riboflavin transporter, whether bacterial (RibU, RibN, RfuABC) (21–23) or eukaryotic (MCH5 in yeast; RFVT and SCL52 familiy members in mammals) (24–27). Mycobacteria have thus been assumed incapable of riboflavin uptake, although the contributions of a non-canonical transporter and/or passive diffusion had not been explored. In our experiments we observed that uptake has an overall linear dependence on riboflavin concentration and does not depend strongly on either proton-motive force or ATP synthesis. This trend is compatible passive diffusion, but does not rule out other possibilities, including facilitated transport mechanisms that are not sensitive to the perturbations tested. It is also possible that a transporter is expressed under other (*e.g.*, stress) conditions. Future studies into the molecular basis of riboflavin uptake in *Mtb* will require additional approaches, such as identification of genes that modulate uptake and biochemical characterization of candidate proteins.

Further, intracellular riboflavin has been reported in the 10-20 µM range in multiple mammalian cell lines (28). This concentration range suggests that *Mtb* could survive in host cells even when riboflavin biosynthesis is inhibited.This raises the question of whether this pathway is a valid drug target, as has been posited and pursued in several Rib enzyme inhibitor development studies (3,4,29–31). However, whether sufficient riboflavin is accessible to *Mtb* or can support growth of a riboflavin auxotroph during infection remained unknown. We therefore asked whether host riboflavin is sufficient *in vivo* using *Mtb* Δ*ribC.* Unexpectedly, we were unable to detect any surviving Δ*ribC in vivo*, as CFU were not recovered from aerosol-infected lungs even at day one post infection and this phenotype was complemented. In contrast, Block et al. reported that *ribA2*::Tn and *ribG*::Tn were recovered and significantly depleted in the lungs at day 21 vs. 1 post infection (12). However, these infections were performed with a library of *Mtb* Tn insertion mutants (105 strains representing 48 genes) and via intravenous route. These key experimental differences could underlie these different observations. If interpreted as a defect in infectivity, our result suggests that loss of riboflavin biosynthesis affects host cell engagement, possibly due to *Mtb* cell surface changes arising indirectly from the shifts in metabolism indicated by our targeted metabolite profiling (**Figure 6, Supplemental Table S5**).

On the other hand, despite significant depletion of intracellular riboflavin after 3 days (**Figure 6A**) and attenuation of growth (**Figure 3C, F**), *Mtb* Δ*ribC* survived for at least 3 weeks in liquid culture following removal of riboflavin from the growth medium. It is possible that micromolar exogenous riboflavin induces non-physiologically high intracellular concentrations that turn over slowly relative to the levels necessary to sustain survival. Thus, the observed weeks-long survival may not accurately represent survival under inhibition of riboflavin biosynthesis. Based on this hypothesis, an inducible knockout that can be propagated in the absence of exogenous riboflavin may help resolve questions around *in vitro* and *in vivo* survival and the validity of riboflavin biosynthesis as a drug target. Indeed, Block, Namugenyi, and Tischler recently reported analogous *in vitro* results in which *ribC*::Tn and *ribA2*::Tn insertion mutants also lost ∼1 log viability over 2 weeks following removal of riboflavin (32). They also blocked *ribA2* transcription using an inducible tet-OFF repressor and found that repression in riboflavin-free medium yielded similar survival loss in the first 2 weeks, but near sterilization after 5 weeks. This inducible system also enabled their observation that *ribA2* repression led to loss of *Mtb* viability in the lungs and spleen at all stages of infection tested.

While targeted metabolomics confirmed reduced riboflavin in *rib* null mutants, other metabolites in the riboflavin pathway such as the intermediates and MAIT ligands 5-OP-RU and DMRL were not found in any strain or sample. This is similar to related studies that have reported MS detection of riboflavin and DMRL, but no other metabolites in the pathway (13,14). In general, the high reactivity of riboflavin and other vitamin intermediates and MR1 ligands pose a significant challenge to their direct detection in biological samples. Future profiling efforts may be aided by stabilization via binding to MR1 (19,33,34). While the relative abundance of these reactive vitamin B metabolites upon *rib* disruption remains unknown, we found an unexpected association between riboflavin biosynthesis and Phe/Tyr metabolism. While there is no obvious direct connection between these two pathways, riboflavin is converted to the cofactors FMN and FAD, which would be required for the function of the approximately 140 predicted flavoproteins in *Mtb* (35). Depletion of riboflavin would therefore be expected to have broad effects via diverse cellular processes. For example, Phe and Tyr accumulation could result from reduced degradation by flavin-containing monoamine oxidase Rv3170 based on KEGG pathways (mtu00350, mtu00360). On the other hand, this effect might be countered by the reduced activity of FMNH_2_-dependent chorismate synthase Rv2540c (36), which feeds into aromatic acid biosynthesis. Another possibility is that disruption of riboflavin production leads to excess precursor ribulose-5-phosphate (**Figure 1**) and thus increased aromatic amino acids by feeding into the non-oxidative pentose phosphate pathway (NOPPP) as well as the shikimate/chorismate pathways (34,35). Overall, disruption of riboflavin biosynthesis could shift the balance among these pathways and thereby contribute to Phe and Tyr accumulation. Beyond amino acids, disrupting riboflavin biosynthesis also affects 6-FP, an inhibitory MR1 ligand and a metabolite derived from folate, whose biosynthesis also depends on chorismate. This is thus also a possible consequence of alterations in flux through shikimate/chorismate pathway and/or FAD/FMN depletion and corresponding loss of function in redox enzymes required for biosynthesis.

In summary our results indicate that following disruption of riboflavin biosynthesis, neither metabolic compensation nor the capacity for riboflavin uptake is sufficient to support infection after murine aerosol infection. Moreover, the broad effects on metabolism suggest that perturbing biosynthesis of one family of MR1 ligands affects the repertoire of MR1 ligands—both activating and inhibitory— produced by *Mtb* as a whole. This interplay may have consequences for our understanding of MR1-mediated immune responses in tuberculosis, as a subject for future studies.

## METHODS

### Bacterial strains and culture conditions

*Mtb* strains H37Rv and mouse-passaged Erdman (gift of Michael Glickman) were cultured in Middlebrook 7H9 medium (HI MEDIA) with 10% v/v oleic acid-albumin-dextrose-catalase (OADC) supplement (BD), 0.5% v/v glycerol and 0.025% v/v Tyloxapol at 37 °C with shaking at 110 rpm. For growth on solid media, *Mtb* were cultured on Middlebrook 7H10 agar with 10% v/v OADC supplement and 0.5% v/v glycerol. Riboflavin was supplemented in medium where noted. Kanamycin (25 μg/mL), hygromycin (50 μg/mL) and zeocin (20 μg/mL) were used as appropriate to select transformants and maintain plasmids.

### CRISPRi mediated knockdown of riboflavin metabolism genes in Mtb

*Mtb* Erdman encoding machinery to knockdown *rv1409* (*ribG*), *rv1412* (*ribC*), *rv1415* (*ribA2*) and *rv1416* (*ribH*) were generated largely as reported (39). Briefly, PAM sequences of varying predicted strengths (low, medium and high) were chosen for each gene from the database available at https://pebble.rockefeller.edu/tools/sgrna-design/. Oligonucleotides complementary to the selected PAM sequences with flanking BsmBI enzyme sites were annealed and ligated into digested pIJR965. Respective PAM sequences and oligonucleotides are listed in **Supplemental Table S2**. The resulting sequence-confirmed vectors as well as pIJR965 as a non-targeting control (NT) were electroporated into *Mtb.* Transformants were selected on solid medium with kanamycin, propagated in liquid medium, and confirmed for knockdown using qRT-PCR as described below. Transformants were also plated for viability on solid media with anhydrotetracycline (ATc, 100 ng/mL).

### Confirmation of CRISPRi knockown with qRT-PCR

Primary cultures for all strains were inoculated from frozen stocks in 7H9 media with kanamycin and cultured for 3-4 days. Once the cultures reached an optical density at 600 nm (OD_600_) of 0.8-1.2, they were subcultured in medium with or without ATc at a starting OD_600_ of 0.25 in 10 mL for 3 days. Bacterial cells harvested by centrifugation at 3000 x*g* at 22 °C were resuspended in 1 mL Trizol LS and lysed by bead beating (Bead Ruptor, Omni Internation) with 0.1 mm zirconia beads for four 30-second cycles with 5 min incubation on ice after each cycle. The suspension was then centrifuged at 10,000 x*g* at 4 °C for 10 min. RNA was extracted from the supernatant by phase separation following addition of chloroform: isoamylalchohol. RNA in the aqueous phase was then isolated (RNeasy Kit, Qiagen) per the manufacturer’s instructions. After DNase digestion with Turbo DNase (Ambion), 1000 ng RNA was used to make cDNA (Verso cDNA synthesis Kit, Thermo Fisher). Purified cDNA was diluted 10-fold and used as template for primer pairs listed in **Supplemental Table S3**. 16srRNA expression was used as housekeeping control.

### Growth and time-kill curves

Primary cultures for all strains were inoculated from frozen stocks in 7H9 medium with kanamycin (CRISPRi knockdown strains) or hygromycin (targeted knockout strains) and cultured for 3-4 days. Once the cultures reached OD_600_ 0.8-1.2, they were subcultured in medium with or without ATc with a starting OD_600_ of 0.015 (CRISPRi knockdown strains) or with and without 20 µM riboflavin, FAD, or FMN supplementation at a starting OD_600_ of 0.1 (targeted knockout strains) in 50 mL in roller bottles. OD_600_ was measured daily. For time-kill curves, after the knockout cultures reached an OD_600_ of 0.8-1.2, they were sub-cultured in media without riboflavin at a starting OD_600_ of 0.01 in 50 mL medium in roller bottles. At indicated time points, appropriate dilutions were plated on 7H10 plates supplemented with riboflavin. CFUs were enumerated after 3-4 weeks.

### Viability spot assays

Logarithmic phase cultures grown as above were diluted to OD_600_ 0.05 and then serially diluted ten-fold up to a factor of 10^5^ dilution. For each dilution 2.5 μL was spotted on medium with or without ATc and varying concentrations of riboflavin as indicated. Plates were imaged after 15-20 days of incubation.

### *Generation of* Δrv1409, Δrv1412, Δrv1415, Δrv1416, *and* Δrv1417

Targeted knockout strains of riboflavin metabolism genes were generated in *Mtb* H37Rv largely as reported (40). Briefly, flanking sequences of 250-500 bp 5’ and 3’ upstream and downstream of the respective genes were cloned on either side of a hygromycin resistance cassette via the HindIII and XbaI sites of pJSC407 (gift of Jeffrey Cox) using InFusion Cloning (Takara Bio). The resulting plasmids (**Supplemental Table S4**) were sequence confirmed and used as a template for PCR with appropriate primers (**Supplemental Table S3**). The resulting purified product was electroporated into *Mtb::*pNIT-RecET expressing the recombinase following induction with isovaleronitrile. Successful recombinants were selected on agar with hygromycin and 20 μM riboflavin. Individual clones were confirmed by PCR and sequencing (**Supplemental Table S3**). Complement constructs were generated by cloning the respective genes and ∼1000 nt 5’ of the start codon (as a native promoter) into a variant of integrating plasmid pMV306 (modified to encode a zeocin resistance cassette) via the XbaI and ClaI sites using InFusion Cloning. The resulting plasmids (**Supplemental Table S4**) were sequence confirmed, electroporated into respective knockout strains, and selected on agar containing zeocin at 10 μg/mL. Strains Δ*rv1415*, Δ*rv1416,* and Δ*rv1417* were generated from Δ*rv1415-rv1416-rv1417* followed by complementation with *rv1416-rv1417*, *rv1415-rv1417,* or Δ*rv1415-rv1416*, respectively (**Supplemental Tables S3, S4**).

### Riboflavin uptake assay

*Mtb* H37Rv (15 mL per condition) were cultured to logarithmic phase (OD_600_ 1-1.2). Following centrifugation, the decanted pellet was resuspended in 1/10^th^ volume of transport buffer (i.e., 1.5 mL of 50 mM KH_2_PO_4_/K_2_HPO_4_, 50 mM MgCl_2_ pH 7.0). ^3^H-riboflavin (Americal Radiolabeled Chemicals or Moravek Chemicals) was added at indicated concentrations followed by incubation at 37 °C for 1 h unless indicated otherwise. Where indicated, carbonyl cyanide 3-chlorophenylhydrazone (CCCP) or bedaquiline (BDQ) at indicated concentrations, or DMSO vehicle control were added and cell suspensions incubated at 37 °C for 30 min before the addition of ^3^H riboflavin. Cell suspensions were then centrifuged at 10,000 xg for 2 min at 22 °C. Pellets were then washed 3 times with 1.5 mL PBS to remove residual ^3^H-riboflavin before resuspension in 1 mL PBS and lysis by bead beating (Bead Ruptor, Omni International) with 0.1 mm zirconia beads for four 30 s cycles with 5 min incubation on ice after each cycle. The suspension was then centrifuged at 10,000 x*g* at 4 °C for 10 min and the supernatant was 0.2-µm filtered to obtain the final clarified lysate. To quantify radioactivity, 200 µL lysate were added to 5 mL scintillation cocktail (Scintiverse) prior to measurement (Beckman Coulter LS6500 Scintillation counter).

### Preparation of metabolite extracts

*Mtb* H37Rv *rib* knockout strains were cultured to logarithmic phase (OD_600_ 1-1.2), centrifuged at 3000 x*g* for 10 min, and resuspended in medium without detergent. 1 mL of the cell suspension was inoculated onto a 0.22 µm PVDF filter (Millipore) as adapted from reference (17) and incubated on 2.7 mL medium with 20 μM riboflavin for 72 h. Filters were then transferred to medium without riboflavin for another 72 h. Filters were transferred into in 2-mL tubes containing 200 µL of 0.1mm zirconia-silica beads and 1 mL 2:2:1 acetonitrile:methanol:water. Cells were lysed by bead beating (Bead Ruptor, Omni International) for six 30-s cycles at 3.25 m/s with a 2-min incubation on ice after each cycle. Filters were removed and the tubes centrifuged at 13,000 x*g* at 4°C for 10 min. The supernatant was then filtered through a nylon 0.2 μm filter (Corning Costar) and further centrifuged at 13,000 x*g* at 4 °C for 5 min. The supernatant was transferred to fresh tubes and submitted for LC-MS analysis.

### Targeted metabolomics by liquid chromatography-mass spectrometry (LC-MS)

For sample extraction, cell extracts were thawed on wet ice for 1 hour. For cell extracts, 10 µL of isotopically labeled riboflavin (^13^C_4_,^15^N_2_-riboflavin, IsoSciences-7072, Entegris; 2 µM stock) was added to 500 µL sample. Cell extracts were dried using a GeneVac concentrator and reconstituted stepwise with 10 µL of acetonitrile:methanol:water (2:2:1, v/v/v), vortexing for 5 s every 5 min for a total of 10 min, followed by the addition of 40 μL of Milli-Q water with vortexing for 5 s every 5 min for an additional 10 min. Reconstituted samples were centrifuged at 20,000 x*g* for 20 min at 4°C and transferred to amber glass autosampler vials.

For LC-MS/MS, samples were analyzed using reverse-phase chromatography on an Agilent 1290 Infinity II LC system coupled to an Agilent 6495C triple quadrupole tandem mass spectrometer (Agilent Technologies), operated in dynamic multiple reaction monitoring (dMRM) with both positive and negative electrospray ionization (ESI) modes. Chromatographic separation was performed on a CORTECS UPLC C18^+^ (2.1 x 150 mm, 1.6 µm, Waters). Mobile phases consisted of (A) 8.7 mM ammonium formate with formic acid (pH 2.9) in water and (B) 100% acetonitrile. The gradient program was as follows: 0-1 min, 0.5% B; 1-12 min, linear increase to 99.5% B; 12-14 min, hold at 99.5% B (0.5 mL/min); followed by 5 minutes of re-equilibration time at initial conditions. The flow rate was 0.4 mL/min; injection volume was 5 µL, and column temperature was maintained at 45°C. Mass spectrometry parameters were as follows: gas temperature, 230°C; gas flow, 11 L/min; sheath gas temperature, 400°C; sheath gas flow, 12 L/min; nebulizer pressure, 25 psi; capillary voltage, +4000 V (positive mode) and -3000 V (negative mode); nozzle voltage, +500 V (positive mode) and -2000 V (negative mode). Data was analyzed using Skyline software, and compound identities were confirmed by comparison to authentic standards.

### Mouse infections

Four-week-old female C57BL/6 mice (C57BL/6NTac, Taconic Biosciences) were housed in the Division of Laboratory Animal Resources for 5 d before being moved to a biosafety level 3 facility where all *Mtb* infections were performed. For infection, cultures of respective strains at OD_600_ 0.8-1.0 were centrifuged at 21 °C at 3,000 x*g* for 5 min and then resuspended in 1XPBS with 0.05% v/v Tween 80. To obtain single-cell suspensions, cultures were sonicated (Q125, Qsonica) with 2 cycles of 5 s on/off at 25% amplitude. After sonication, OD_600_ was measured again and a final inoculum of 30 ml for infection was made in sterile water at a concentration of 3.6 x 10^4^ CFU/mL (using the conversation OD_600_ 1 = 3 x 10^8^ CFU) for low dose infection. Each mouse was loaded in a nose-only restrain tube (CH Technologies), with a circular pusher with nose only exposed to the vent on a Biaera Aero3G (Biaera Technologies). The aerosol infections with *Mtb* strains were carried out using Aero3G v7.60.1 software with the following configuration: Chamber: ONARES; flow: 15.00 lpm; generator: 3-Jet collision with flow of 7.50 lpm; time of exposure: 10 min; purge: 5 min. Environmental conditions for the instrument were 24°C with humidity 42%. At days 1, 7, and 14 post infection, mice were sacrificed by CO_2_ exposure. Lungs were homogenized in 1 ml PBS with 0.05% Tween 80, ∼10 zirconium oxide beads (2.0 mm), and subjected to bead beating for 5 min at speed 16 (Bullet Blender 5E Pro, Next Advance). Homogenized lung suspensions were serially diluted in PBS with 0.05% Tween 80 and plated on 7H10 agar. Plates were incubated at 37 °C, with 5% CO_2_, and colonies were enumerated after 3 weeks.

## ACKNOWLEDGMENTS

The authors acknowledge support from Stony Brook Foundation, the Stony Brook University Office of the Vice President for Research and Department of Medicine of the Renaissance School of Medicine (all to C.K.V.). We also acknowledge support to A.A. from T32 GM127253 and from R21 AI183259 (J.C.S.), R21 AI171579 (C.K.V.) and a Potts Memorial Foundation Award (C.K.V.).

