## Supplemental Figures S1-S5 for "Loss of riboflavin biosynthesis leads to accumulation of select aromatic amino acids and loss of infectivity in *Mycobacterium tuberculosis*"

**Supplemental Material**

- **Figure S1.** Encoding sgRNAs for CRISPRi knockdown of *ribG* does not result in decreased RNA levels upon ATc induction.
- **Figure S2.** *rib* knockdown strains induced with ATc in the absence of riboflavin recover growth after culturing without kanamycin selection.
- **Figure S3.** Riboflavin is significantly depleted only in the *ribA2* CRISPRi knockdown strain at 4 days following ATc induction.
- **Figure S4.** qRT-PCR confirms changes in gene expression in corresponding targeted *rib* null and complement strains.
- **Figure S5.** Riboflavin supplementation does not affect the growth of wild-type or *rib* complement strains.
- **Figure S6.** Neither FAD nor FMN rescues the growth of *rib* targeted null strains.

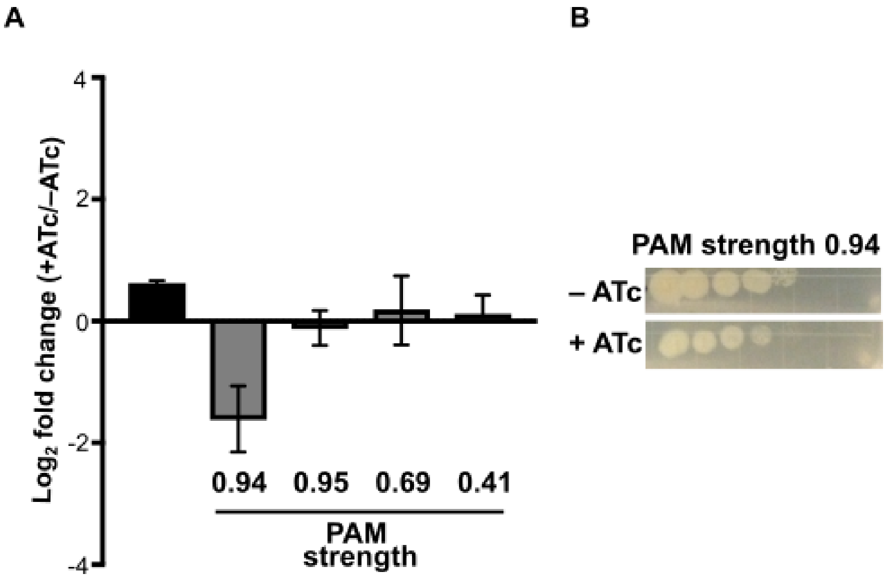


**Figure S1. Encoding sgRNAs for CRISPRi knockdown of *ribG* does not result in decreased RNA levels upon ATc induction. (A)** Erdman *Mtb* strains encoding sgRNA with varying PAM strengths targeting *ribG* (grey bars) or a non-targeting (NT) control (black bar) were cultured with 0 or 100 ng/mL ATc for 3 days. Total RNA extracted from each condition was used to generate cDNA for qRT-PCR. Raw Ct values obtained for respective genes in each condition were normalized to the housekeeping gene *16SrRNA* and the ratio +ATc/−ATc calculated to determine fold change. No changes were statistically significant by unpaired Student's *t*-test. Data shown are the mean ± SEM from 3 independent experiments. **(B)** Strains grown as in (A) were diluted to $10^5$ cells/ml and spotted in serial 10-fold dilution on agar with or without 100 ng/mL ATc.

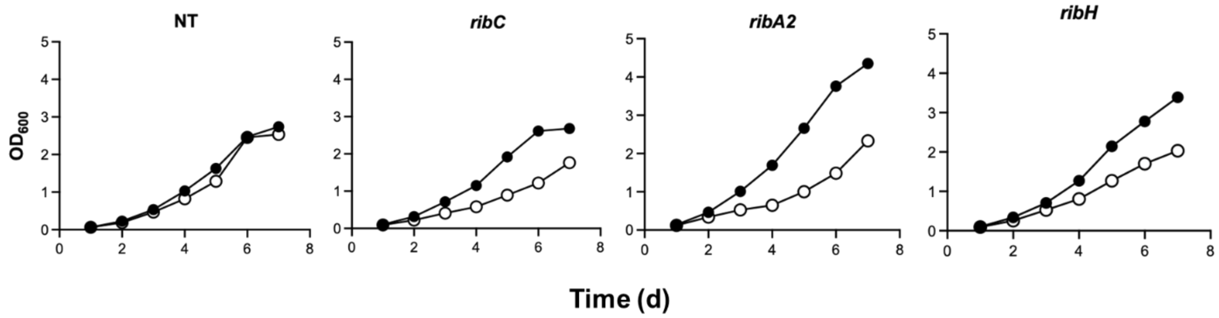


**Figure S2. *rib* knockdown strains induced with ATc in the absence of riboflavin recover growth after culturing without kanamycin selection.** Erdman *Mtb* strains encoding sgRNA for **(A)** a non-targeting (NT) control, **(B)** *ribC*, **(C)** *ribA2*, or **(D)** *ribH* were grown to log phase and then subcultured to $OD_{600}$ 0.015 in 7H9 medium without kanamycin and with 0 (black) or 100 (white) ng/mL ATc. $OD_{600}$ was measured daily for 7 d. Data shown are from one independent experiment.

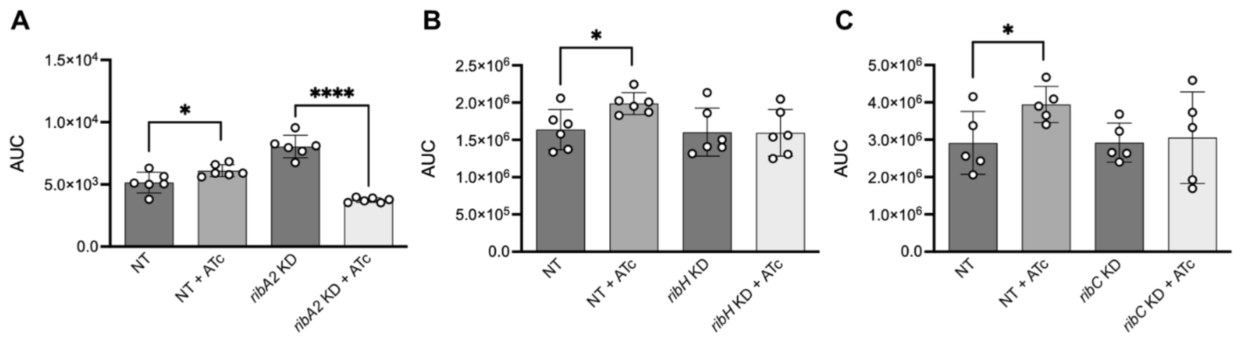


**Figure S3. Riboflavin is significantly depleted only in the *ribA2* CRISPRi knockdown strain at 3 days following ATc induction.** Intracellular riboflavin was quantified by LC-MS from extracts of Erdman Mtb strains encoding CRISPRi sgRNA targeting strains as follows: **(A)** *Mtb* Erdman non-targeting (NT) control and *ribA2* knockdown +/- 100 ng/mL ATc for 3 d, **(B)** *Mtb* Erdman NT and *ribH* knockdown +/- 750 ng/mL ATc for 3 d, and (C) Mtb Erdman NT and *ribC* knockdown +/- 750 ng/mL ATc for 6 days. Data shown are mean area under the curve (AUC) for riboflavin, n = 5-6 biological samples. Samples for each knockdown strain and a NT comparator were prepared together in one independent experiment. Statistical significance was determined by unpaired Student's t-test (*$p$ <0.05, ****$p$ <0.0001).

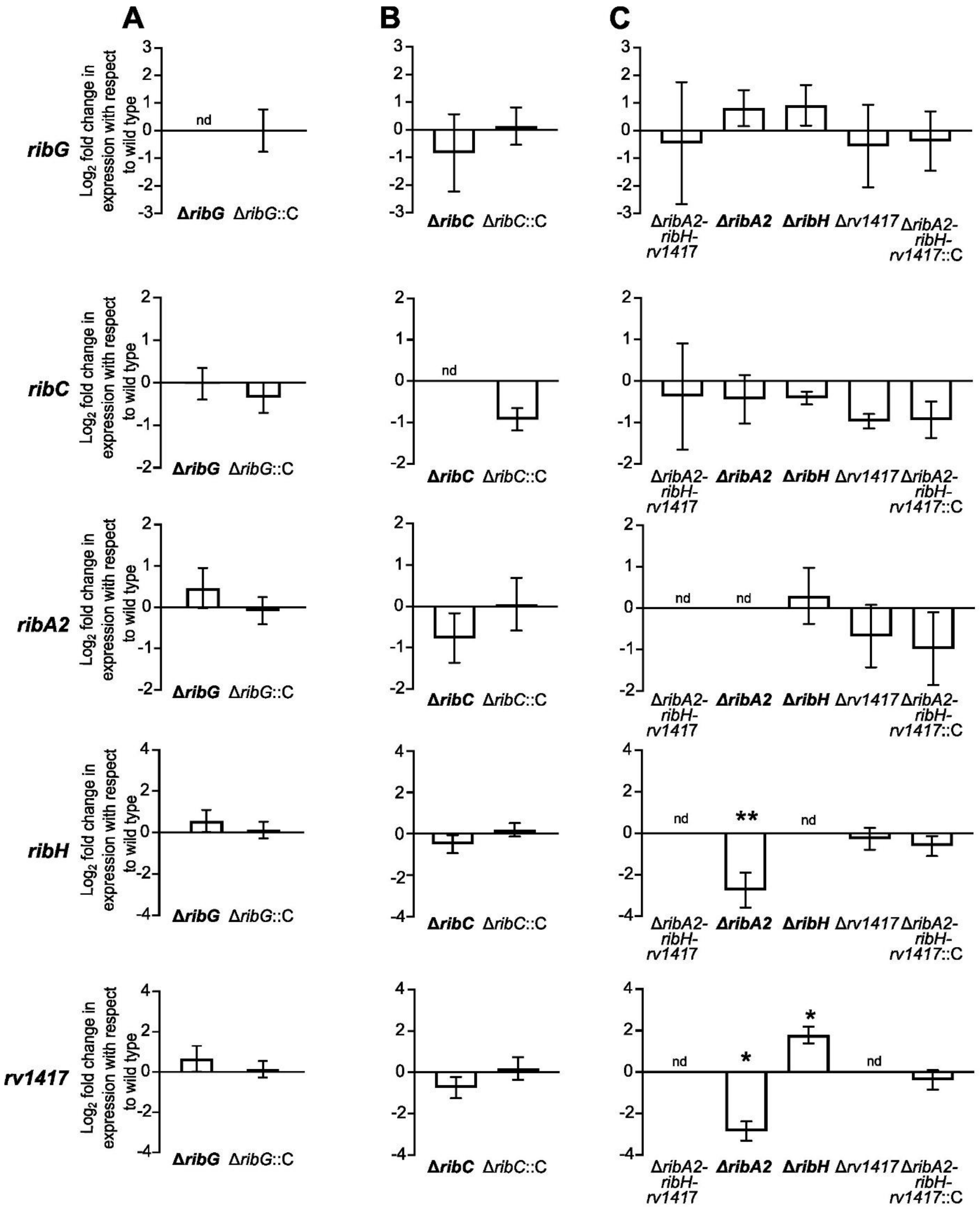


**Figure S4. qRT-PCR confirms changes in gene expression in corresponding targeted rib null and complement strains.** *Mtb* H37Rv wild type and corresponding complements for **(A)** Δ*ribG*, **(B)** Δ*ribC*, **(C)** the operon knockout Δ*ribA2-ribH-rv1417* and derivative strains Δ*ribA2*, Δ*ribH,* Δ*rv1417* were grown in the presence of 20 μM riboflavin to $OD_{600}$ 1. Total RNA extracted from each sample was used to generate cDNA for qRT-PCR. Raw Ct values obtained for respective genes in each condition were normalized to the housekeeping gene *16SrRNA* and the ratio knockout/wild type calculated to determine fold change. Data shown are mean ± SEM from 3 independent experiments. Statistical significance for knockout vs. wild type was determined by unpaired Student's t-test (* $p$ <0.05, ** $p$ <0.01).

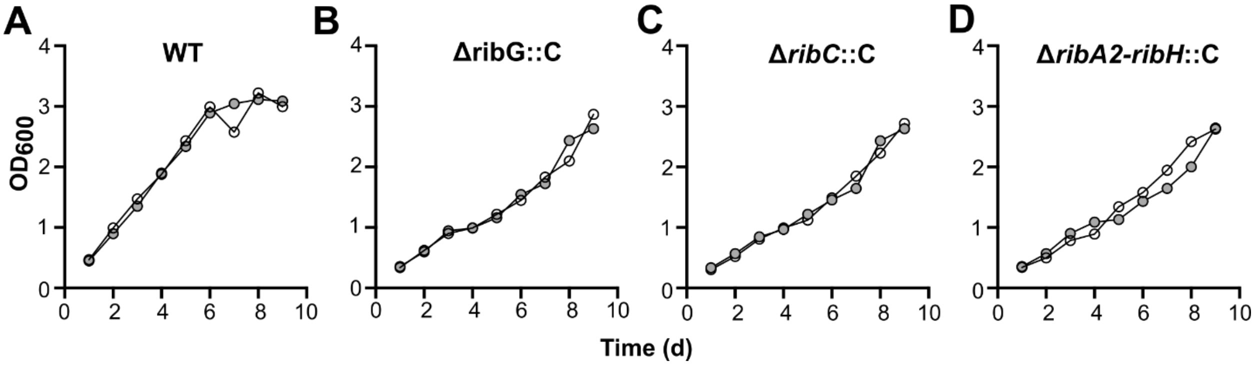


**Figure S5. Riboflavin supplementation does not affect the growth of wild-type or *rib* complement strains.** Growth of *Mtb* H37Rv **(A)** wild type and the complement strains (abbreviated as “C”) **(B)** Δ*ribG::ribG,* **(C)** Δ*ribC::ribC* and **(D)** Δ*ribA2-ribH-r1417*::*ribA2-ribH-rv1417* (operon complement) as measured by $OD_{600}$ daily for 9 d without riboflavin (white circles) or with 20 µM riboflavin (grey circles). Data are from one experiment.

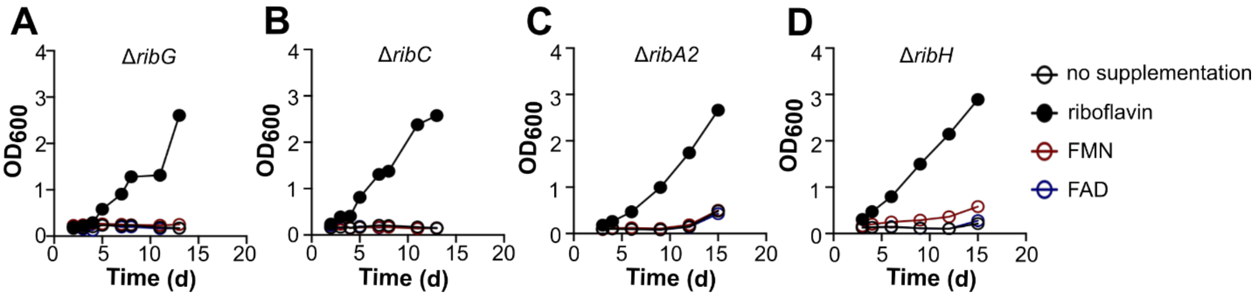


**Figure S6. Neither FAD nor FMN rescues the growth of targeted *rib* knockout strains.** Growth of *Mtb* H37Rv strains **(A)** Δ*ribG,* **(B)** Δ *ribC*, **(C)** Δ*ribA2*, and **(D)** Δ*ribH* as measured by $OD_{600}$ daily for 12 d without riboflavin (white circles) or with 20 µM riboflavin (black circles), FMN (red circles) or FAD (blue circles). Data are from one experiment.
